# KCNK18 c.1107del frameshift in peripheral neuropathy links TRESK dysfunction to neuropathic pain

**DOI:** 10.64898/2025.12.08.692748

**Authors:** Clara Rubio, Milena Ślęczkowska, Enric Pisa, Andrea Barbieri, Anna Pérez, Gerard Callejo, Margherita Marchi, Erika Salvi, Grazia Devigili, Janneke GJ Hoeijmakers, Catharina G Faber, Monique M Gerrits, Hubert JM Smeets, Guillaume Sandoz, Xavier Gasull, Núria Comes

## Abstract

Neuropathic pain is commonly accompanied by hyperexcitability of nociceptive neurons, which is driven by regulation of several ion channels, including two-pore domain potassium channels (K2P) that stabilize the resting membrane potential. The TWIK-Related Spinal cord potassium channel (TRESK, K2p18.1) is predominantly expressed in sensory ganglia, where it provides major background K⁺ conductance. We describe a heterozygous c.1107del frameshift mutation in *KCNK18*, encoding TRESK, found in a patient with painful small fiber neuropathy and dysautonomia. This C-terminal c.1107del mutant is predicted to generate an elongated C-terminal domain, with an altered protein sequence. Whole-cell patch calmp recordings showed that the c.1107del variant reduced K⁺ current density, while heterozygous-like expression resulted in intermediate currents, consistent with a haploinsufficiency mechanism. Confocal imaging revealed decreased plasma membrane expression, indicating a trafficking-dependent loss of function while oligomerization with TRESK (homomeric channels) or TREK1 remained intact. In silico analysis revealed an altered phosphorylation pattern and reduced C-terminal hydrophobicity in the mutant TRESK. Overall, these findings support c.1107del as a pathogenic *KCNK18* variant causing TRESK haploinsufficiency, offering mechanistic insight into the pathophysiology of neuropathic pain.

## Introduction

The precise regulation of primary sensory neuron excitability relies on the coordinated activity of ion channels that govern membrane potential and action potential firing. Among these, background potassium channels of the two-pore domain (K2P) family are important contributors to resting membrane potential (RMP) and play a critical role in the subthreshold range of the membrane potential, where most other conductances are inactive. Therefore, their dysregulation can lead to hyperexcitability underlying diverse neurological disorders. Aberrant sensory neuron excitability is a hallmark of neuropathic pain, a debilitating condition driven by maladaptive changes in neurons of the dorsal root and trigeminal ganglia^1^. In small and medium-sized nociceptors, the K2P TWIK-Related Spinal cord K⁺ (TRESK) channel, encoded by *KCNK18*^2–4^, provides a prevalent background K⁺ conductance that stabilizes the resting membrane potential, shape nociceptor excitability and modulates pain signaling^2,5,6^.

Consistently, TRESK knock-out mice manifest an increased mechanical sensitivity and cold allodynia without affecting thermal sensitivity^7^. Likewise, reduction or inhibition of TRESK heightens nociceptor excitability, as seen in DRG neurons from the G339R *KCNK18* knockout mice^2^, after sciatic nerve axotomy^8^, CFA-induced inflammation^9^ and in cancer pain models^10^. Intrathecal TRESK-siRNA reduces mechanical pain thresholds without affecting thermal sensitivity^11^, resembling nerve injury-induced hypersensitivity^8^. Similarly, pharmacological inhibition of TRESK induces nocifensive behaviors in healthy rats^8^ while its overexpression in DRG neurons alleviates mechanical allodynia and neuropathic pain^12,13^. Reduction of TRESK expression reveal increased DRG neuron excitability and behavioral hypersensitivity to noxious and non-noxious stimuli, while RMP remains unchanged^2,7,14^, likely due to compensatory activity of other channels. Together, these findings highlight the crucial role of TRESK in nociceptor excitability, provide mechanistic insight into neuropathic persistent pain, and suggest that it may represent a promising target for analgesic intervention.

Chronic pain disorders are a multifaceted challenge with highly complex etiology and critically limited therapeutic options. One illustrative example is neuropathic pain (NeuP), which can result from metabolic disorders, infections, chemotherapy, and trauma, often requiring a multidisciplinary approach^12,15^. Small-fiber neuropathy (SFN), a subtype of NeuP, involves damage to unmyelinated C fibers and thinly myelinated Aδ fibers, causing symptoms including spontaneous stabbing or burning pain, itching, numbness, or allodynia^1,16^. Managing NeuP remains particularly difficult due to limited treatments, adverse effects, and variable responses^17,18^. It extends far beyond physical symptoms, impairing emotional, social and economic functioning and stability. This cumulative challenge makes this population especially vulnerable, reinforcing the urgency of identifying the molecular mechanisms underlying neuronal hyperexcitability to develop more effective, targeted therapies.

Pain genetics research has largely focused on the SCN family, encoding voltage-gated sodium channels (Nav) and crucial for action potential generation in nociceptors. Gain-of-function (GoF) mutations in SCN9A (Nav1.7) increase excitability and spontaneous nociceptor firing in various pain disorders^19–21^, while SCN11A (Nav1.9) GoF variants have been associated with painful peripheral neuropathy^22^ and familial episodic pain^23^. In contrast, the link between *KCNK18* and altered nociception remains rarely reported^24–26^. The F139WfsX24 frameshift mutation, associated with familial autosomal dominant migraine with aura, abolishes background currents by preventing membrane trafficking. It also forms heteromeric complexes and exerts a dominant-negative effect on TREK1/2, driving trigeminal neuron hyperexcitability. Remarkably, its correction by CRISPR-Cas9 reverses the sensory neuron hyperexcitability that may contribute to migraine pain induction^27–29^. Beyond migraine, the p.Tyr163Asp and p.Ser252Leu variants associated with intellectual disability and autism impair channel activity via calcineurin-dependent mechanisms.^30,31^. These findings offer insight into how altered TRESK may influence complex disorders associated with neuronal excitability.

In this study, we characterize the *KCNK18* c.1107del frameshift mutation in the C-terminal domain of TRESK, identified in a patient with small fiber neuropathy. This variant provides a valuable model to uncover how TRESK dysfunction may contribute to chronic pain mechanisms^32,33^. We propose that p.(Met370Cysfs) cause TRESK dysfunction, which will lead to nociceptor hyperexcitability, contributing to the patient’s chronic pain. The primary aim of this study is to functionally characterize the c.1107del variant, evaluating its impact on TRESK activity and providing potential mechanisms underlying neuropathic pain pathophysiology.

## Methods

### Plasmid Constructs

Human TRESK pcDNA3.1(+) vector was kindly provided by Dr. Y. Sano (Astellas Pharma Inc, Ibaraki, Japan) and subcloned into pEGFP-C2 (Clontech) vector with an *EcoRI/SmaI* digest (pEGFP-hTRESK)^34^. pEGFP-mTREK-1 was a kind gift from Dr. G. Sandoz (CNRS-Universite de Nice-Sophia Antipolis, France). Additionally, the plasmid was digested with ApaI and EcoRI-HF, and the excised *KCNK18* coding sequence was subcloned into mCherry-C2 using T4 DNA ligase (mCherry-hTRESK). The c.1107del mutation was introduced using the QuikChange II XL site-directed mutagenesis kit (Agilent Technologies, Santa Clara, CA) with primers designed using the QuikChange Primer Design Tool (Forward 5’ GCTGATTGACATATACAAAAATGTATGCTATTCTTTGCAAAAGGGAAG-3’, Reverse 5’-CTTCCCTTTTGCAAAGAATAGCATACATTTTTGTATATGTCAATCAGC-3’).

Mutant plasmids were transformed into XL10-Gold ultracompetent cells, amplified using standard PCR, and verified by Sanger sequencing (Stabvida, Caparica, Portugal). Expression plasmids were then transformed into E. coli DH5α competent cells using the NucleoSpin kit (Macherey-Nagel GmbH & Co. KG, Düren, Germany). Sequences were analyzed using APE v2.0.55 (A Plasmid Editor; M. Wayne Davis, University of Utah). Constructs for functional studies included WT pEGFP-hTRESK, mCherry-hTRESK and pEGFP-hTRESK-c.1107del.

### Cell culture and transfection

Cultured human embryonic kidney (HEK-293T) cells, obtained from the American Type Culture Collection (AITCC), were maintained in Dulbecco’s Modified Eagle Medium (DMEM) supplemented with 1 mg/mL fetal bovine serum (FBS), 100 mg/mL of penicillin / streptomycin and 100 mg/mL of L-glutamine (Sigma-Aldrich, Madrid, Spain) at 37°C in humidified atmosphere with 5% CO_2_. Upon reaching confluence, cells were passed using trypLE™ express (Invitrogen, Thermo Fisher Scientific, Carlsbad, CA). To study homomeric channels, transient transfections with 500 ng of either WT pEGFP-hTRESK or pEGFP-hTRESK-c.1107del were performed using RotiFect™ (Carl ROTH GmbH, Karlsruhe, Germany) 24 hours prior to electrophysiological recording, confocal microscopy or oligomerization analysis. In some studies, cells were co-transfected with 250 ng of mCherry-hTRESK and 250ng of pEGFP-hTRESK-c.1107del to mimic the patient’s heterozygous condition.

### Electrophysiology

Electrophysiological recordings were performed with a patch-clamp amplifier (Axopatch 200B, Molecular Devices, LLC, San Jose, CA), an Axiovert 35M inverted phase contrast microscope (Zeiss, Jena, Germany), and a PatchStar micromanipulator (Scientifica Ltd., Uckfield, UK). Patch electrodes (Warner Instruments, Hamden, CT) were fabricated in a Flaming/Brown micropipette puller P-97 (Sutter instruments, Novato, CA). Electrodes had a resistance between 2 and 4 MΩ when filled with intracellular solution (in mM): 135 KCl, 2 MgCl_2_, 2.1 CaCl_2_, 2.5 Na_2_-ATP, 5 EGTA, 10 HEPES, pH adjusted to 7.25 with KOH (osmolality 305.5 ± 2.8 mOsm kg^−1^). Bath solution contained (in mM): 145 NaCl, 5 KCl, 2 CaCl_2_, 2 MgCl_2_, 10 HEPES-NaOH, 10 Glucose, pH adjusted to 7.4 with NaOH (osmolality 296.6 ± 1.9 mOsm kg^−1^). Membrane currents were recorded using the whole-cell patch-clamp configuration, filtered at 2 kHz and digitized at 10 kHz. Series resistance was kept below 15 MΩ and compensated at 70–80%. Current density (pA/pF) was calculated by dividing the measured current amplitude at a given voltage pulse by the cell’s membrane capacitance. All recordings were made at room temperature (22-23°C). Data were analyzed using pCLAMP 10.6 (Molecular Devices) and GraphPad Prism 10 (GraphPad Software, Inc., La Jolla, CA). TRESK currents were assessed using either voltage ramps or voltage step pulses lasting 400 msec. The holding potential was set to −60 mV and voltage ramps ranged from −100 mV to +50 mV, while step pulses were applied from −100 mV to +60 mV. The reversal potential (E_rev_) was obtained for each cell from the intersection of the I-V curve. The apparent conductance was calculated as G(V)=I(V)/(V−E_rev_) and normalized to G_max_. Points close to E_rev_ were excluded to avoid numerical instabilities. The normalized curves were fitted with a standard Boltzmann equation G∼(V)=offset+amplitude/(1+exp(−(V_1/2_−V)/slope)); with the slope being RT/zF (R, universal gas constant; T, temperature; z, equivalent gating charge [e_0_]; F, Faraday constant) to estimate V_1/2_ and the slope (k).

### Single Molecule Pulldown

SiMPull experiments were done as previously described^29^. A DNA ratio of 1:1 was used. Constructs used were pEGFP-hTRESK, pEGFP-hTRESK-c.1107del, HA-hTRESK-pCDNA3.1, HA-TREK1-pCDNA3.1. Briefly, 24 hours after transfection, HEK 239T cells were harvested from coverslips with Ca^2+^-free PBS buffer and lysed in buffer containing (in mM): 150 NaCl, 10 Tris pH 7.5, 1 % EDTA, protease inhibitor cocktail (Thermo Scientific) and 1.5% IGEPAL (Sigma) or 1% DDM (Sigma). After 30-60 minute incubation at 4°, lysate was centrifuged and the supernatant was collected. Coverslips passivated with PEG (∼99%)/ biotin-PEG (∼1%) and treated with NeutrAvidin (Pierce) were used. 15 nM biotinylated anti-HA antibody (clone 16B12, BioLegend) was applied for 20 minutes and then washed out. Antibody dilutions and washes were done in T50 buffer with BSA containing (in mM): 50 NaCl, 10 Tris pH 7.5, and 0.1 mg/mL BSA. Lysate, diluted in lysis buffer containing 0.04% IGEPAL, was then applied to the chamber and washed away following brief incubation (∼2 minutes). Single molecules were imaged using a 488 nm Argon laser on a total internal reflection fluorescence microscope with a 60x objective (Olympus). We recorded the emission light after an additional 3x magnification and passage through a double dichroic mirror and an emission filter (525/50 for GFP) with a back-illuminated EMCCD camera (Andor iXon DV-897 BV). Movies of 250-500 frames were acquired at frame rates of 10–30 Hz. The imaged area was 13 x 13 μm2. At least 5 movies were recorded for each condition and data was analyzed using custom software. Multiple independent experiments were performed for each condition. Representative data sets are presented to quantitatively compare conditions tested on the same day.

### Colocalization analysis by confocal microscopy

Cells were seeded on poly-L-lisine-coated glass coverslips and transfected 24 h prior to imaging. Plasma membrane were stained with Concanavalin A-Alexa 594 at 5 µg/mL in PBS for 10 min at 4 °C, followed by 3x PBS washes and by fixation with 4% paraformaldehyde in PBS for 10 min at RT. Samples were mounted with ProLong™ Gold Antifade Mountant containing DAPI (Thermo Fisher Scientific, Waltham, MA) and protected from light until imaging. Confocal images were acquired on a Leica TCS SP5 laser-scanning confocal microscope using a 40× oil-immersion objective lens and the Leica Application Suite AF software (Leica Microsystems). Images were collected at a resolution of 1024x1024 pixels, 16-bit depth, using sequential scanning to avoid spectral overlap. Excitation/emission settings were: DAPI (nuclei), UV/420–480 nm; EGFP (c.1107del), 488 nm/500–550 nm; mCherry (WT), 561 nm / 600–670 nm; and Concanavalin A-Alexa 594 (membrane marker), 594 nm / 600–630 nm. Offline image analysis was done using CellProfiler pipelines. Nuclei (DAPI) and plasma membrane regions of interest (ROIs) were automatically segmented, and mean fluorescence intensities for WT and c.1107del were measured within membrane and cytoplasmic compartments. Colocalization was quantified using Mander’s overlap coefficient, to measure the fraction of one fluorophore’s intensity overlapping with another. Values range from 0 (no overlap) to 1 (entire colocalization).

### Data analysis

Data are presented as mean ± SEM. Statistical differences between different sets of data were assessed by performing unpaired Student’s t-tests or one-way ANOVA followed by Šidák multiple comparisons test. The significance level was set at P<0.05 in all statistical analyses. Data analysis was performed using GraphPad Prism 10 software (GraphPad Software, Inc.).

## Results

### Clinical features and predicted structural impact of the c.1107del mutation

A screening of peripheral ion channel genes in a cohort of patients with painful small fiber neuropathy (SFN) at Maastricht University Medical Centre+ (all tested negative for pathogenic variants in major voltage-gated sodium channels; SCN genes), identified a heterozygous single-nucleotide deletion at position 1107 (c.1107del) in the *KCNK18* gene, which encodes for the TRESK potassium channel (K2p18.1). This variant was found in a 60-year-old woman with a three-year history of progressive SFN characterized by burning and tingling pain, distal paresthesia, and dysautonomia including hot flashes, dry eyes, constipation and increased transpiration. Standard analgesics provided little relief, and the absence of a family history suggests a sporadic occurrence^32,33^. According to the American College of Medical Genetics and Genomics (ACMG) guidelines^35^, c.1107del is classified as a variant of uncertain significance (VUS). The single-thymine deletion occurs within the distal coding region of TRESK and introduces a frameshift starting at codon Met370, abolishing the canonical stop codon and redirecting translation into a novel out-of-frame sequence until a downstream termination site is reached. As a result, the mutant protein p.(Met370Cysfs) is predicted to replace the final 29 C-terminal residues of the wild-type channel with a novel extension spanning 384 to 397 (Fig. 1). This altered C-terminus provides a plausible molecular basis for the patient’s clinical phenotype, although its precise structural consequences remain to be experimentally validated.

**Fig 1.**
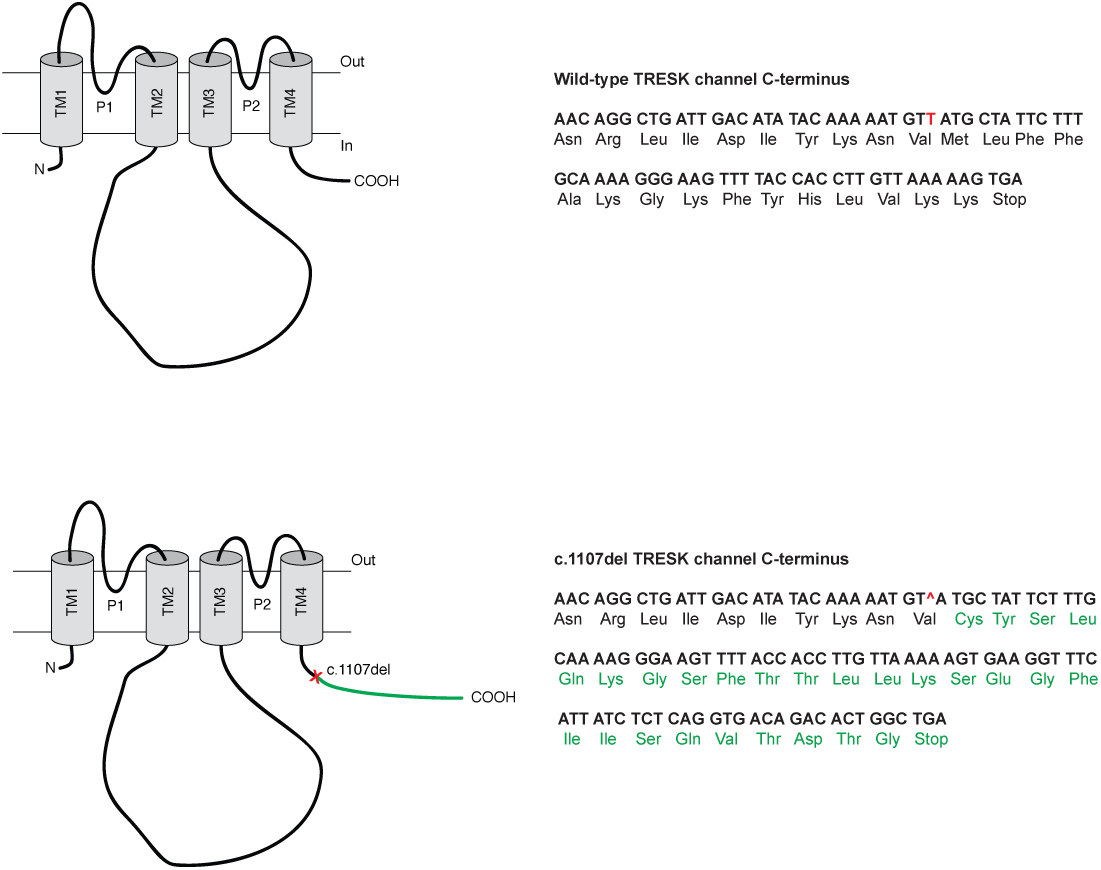
Schematic representation of wild-type and c.1107del TRESK, with sequences of the distal C-terminal region. The c.1107del *KCNK18* variant causes a frameshift, resulting in a predicted extension of the C-terminal domain and a new amino acid sequence highlighted in green.

### The c.1107del *KCNK18* variant reduces TRESK-mediated potassium currents

To determine the functional impact of the patient’s c.1107del genetic variant, we performed whole-cell patch-clamp recordings in HEK-293T cells expressing wild type (WT) or mutant TRESK. Representative whole-cell currents evoked by a voltage-step protocol were recorded from cells expressing either WT hTRESK or hTRESK-c.1107del plasmids (Fig. 2A). WT TRESK currents exhibited the characteristic outwardly rectifying I-V relationship typical of this channel in a physiological K^+^ gradient^3,4^. Mean amplitudes were −550.9 ± 71.0 pA at −100 mV and 4254.7 ± 421.4 pA at +50 mV (n=22). In contrast, the c.1107del mutant channel displayed markedly smaller currents of −250.1 ± 41.4 pA at −100 mV and 1833.0 ± 150.6 pA at +50 mV (P ≤ 0.0001 one-way ANOVA, n=41; Fig. 2B), corresponding to a reduction of 55% and 52%, respectively. Current traces obtained with voltage ramps yielded comparable I-V curves to those obtained with voltage pulses. Therefore, for consistency and to streamline data presentation, subsequent analyses were based on voltage ramp-evoked currents, from which current densities (pA/pF) were calculated.

**Fig 2.**
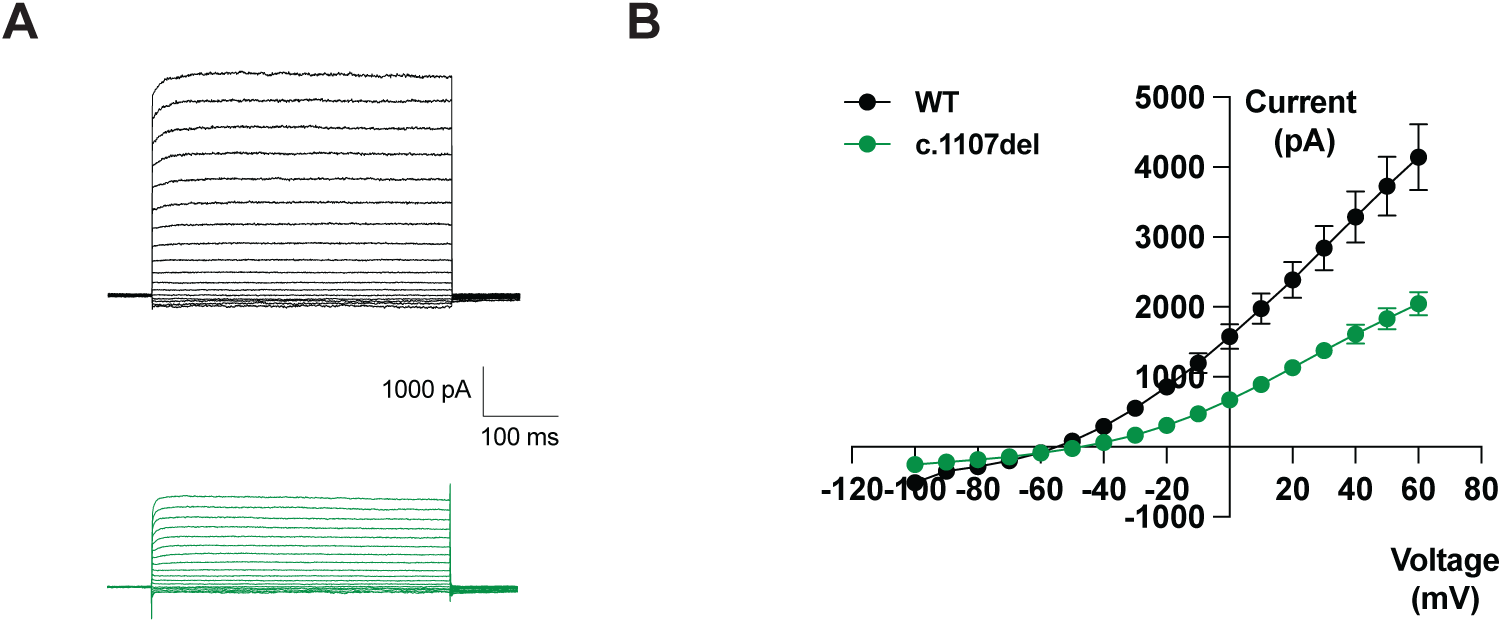
Effect of c.1107del on TRESK-mediated current amplitudes. **A.** Representative whole-cell patch clamp current recordings from cells expressing wild-type (WT; grey) or c.1107del (green) TRESK. Currents were elicited using a voltage-ramp protocol (from –100 mV to +50 mV), and the resulting traces illustrate the reduced amplitude in the c.1107del currents compared with WT. **B.** Current-voltage (I-V) relationships are plotted from voltage pulses and illustrate reduced currents across the voltage range for the c.1107del. Values represent mean ± SEM (n=22 for WT, n=41 for c.1107del, P≤0.0001 one-way ANOVA).

When normalized to cell capacitance to account for differences in cell size, current densities confirmed the decrease being −17.8 ± 2.3 pA/pF for WT vs. −8.4 ± 0.8 pA/pF for c.1107del at −100 mV, and 175.2 ± 24.7 pA/pF vs. 82.5 ± 6.3 pA/pF at +50 mV (P < 0.0001, one-way ANOVA). These values represent a 53% reduction in current density in c.1107del mutants (n=12) compared to WT (n=24) at both −100 mV and +50 mV (Fig. 3B). To mimic the heterozygous state identified in the patient, HEK-293T cells were co-transfected with equal amounts of WT and c.1107del constructs (1:1 ratio). Whole-cell recordings under these heterozygous conditions revealed intermediate current densities between those observed in WT and mutant in homozygous conditions (−11.2 ± 0.5 pA/pF at −100 mV and 112.7 ± 10.1 pA/pF at +50 mV), representing reductions of 37% and 36% compated to WT-expressing cells, respectively (n=12, P=0.038 and P=0.041, one-way ANOVA; Fig. 3B). These intermediate values suggest a partial loss-of-function consistent with TRESK haploinsufficiency. Cell capacitance did not differ significantly across conditions (WT: 26 ± 2.8 pF, n=26; c.1107del: 26 ± 2.0 pF, n=45; heterozygous: 23 ± 1.2 pF, n=12), confirming the validity of the calculated current densities (data not shown).

**Fig 3.**
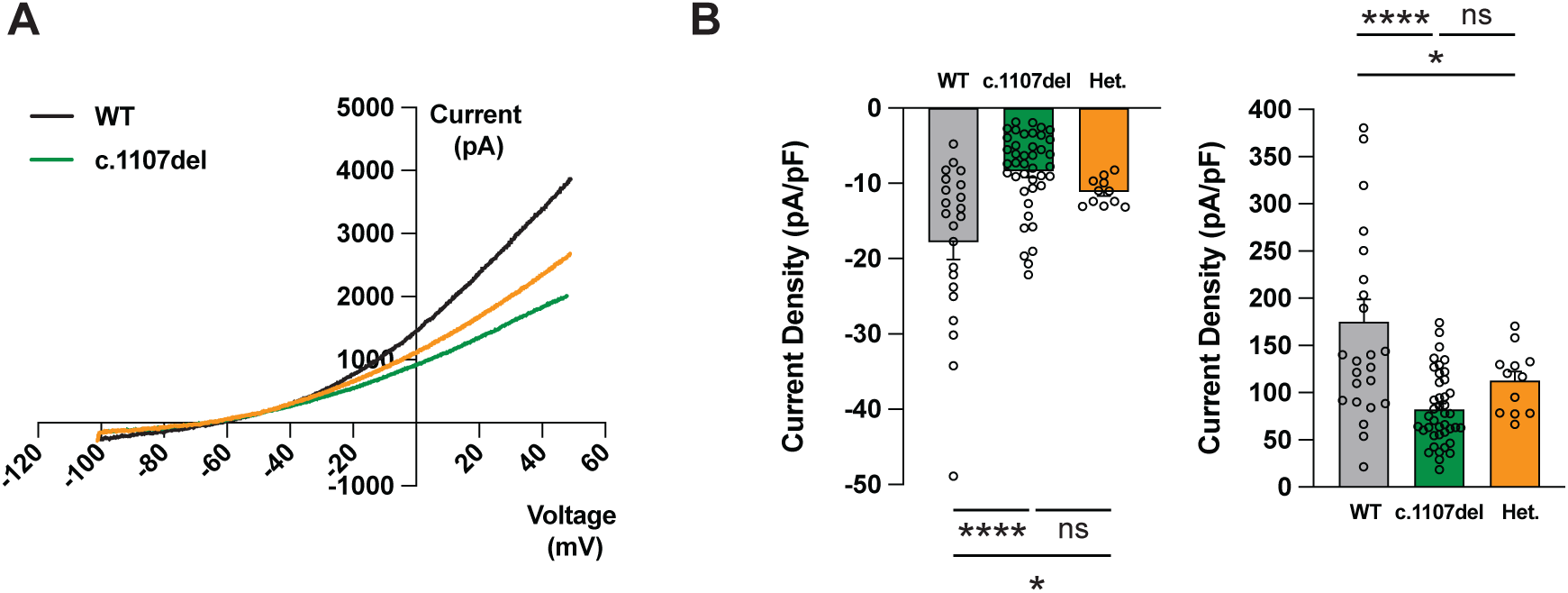
Reduced TRESK current density in c.1107del mutant and heterozygous channels. **A.** Current amplitudes were quantified from whole-cell recordings in cells expressing WT (grey), c.1107del (green), or heterozygous (WT+c.1107del, orange) TRESK. **B.** Bar plots show current densities (pA/pF) measured at –100 mV and +50 mV from voltage-ramp protocols. c.1107del displays significant reduced current densities at both voltages compared with WT, while heterozygous cells exhibit intermediate values. Each point represents an individual cell; bars show mean ± SEM (n=22-24 for WT, n=41 for c.1107del, n=11-12 for WT+c.1107del, ns non-significant P>0.05, *P < 0.05, ****P≤0.0001, one-way ANOVA plus Šídák’s post-tests).

Altough the change in the C-terminal domain of the channel did not predicted an alteration of the biophysical properties of the channel, we evaluated the ionic selectivity by recording TRESK-mediated currents under physiological (5 mM) and elevated (30 mM) extracellular potassium conditions. As predicted by the Nernst equation, the reversal potential (E_rev_) of TRESK shifted from −63.8 ± 3.1 mV (at 5 mM [K⁺]_e_) to −40.8 ± 2.7 mV (at 30 mM [K⁺]_e_), indicating a K⁺-selective conductance. Similar shifts were obtained in cells expressing the c.1107del mutant channel, with reversal potentials of −60.3 ± 1.7 mV (5 mM [K⁺]_e_) and −39.6 ± 0.9 mV (30 mM [K⁺]_e_), confirming that the mutant preserved the typical K⁺-selectivity of TRESK (n=7, Fig. 4A). Accordingly, the direction of K⁺ flux reversed around the calculated equilibrium potential, with significantly greater inward currents measured at −90 mV under high [K⁺]_e_ than outward currents at +45 mV (n=14-22). These findings are consistent with a marked increase in K⁺ influx together with a reduced driving force for K⁺ efflux, supporting the K⁺ selectivity of TRESK and the expected shift in electrochemical equilibrium induced by elevated [K⁺]_e_. As observed for WT, cells expressing c.1107del displayed larger inward than outward currents under high [K⁺]_e_, consistent with preserved K⁺ selectivity. These results further indicate that the mutant channel retains the characteristic K⁺-selective permeation profile of TRESK (n=13, unpaired Student’s t-test; Fig. 4B).

**Fig 4.**
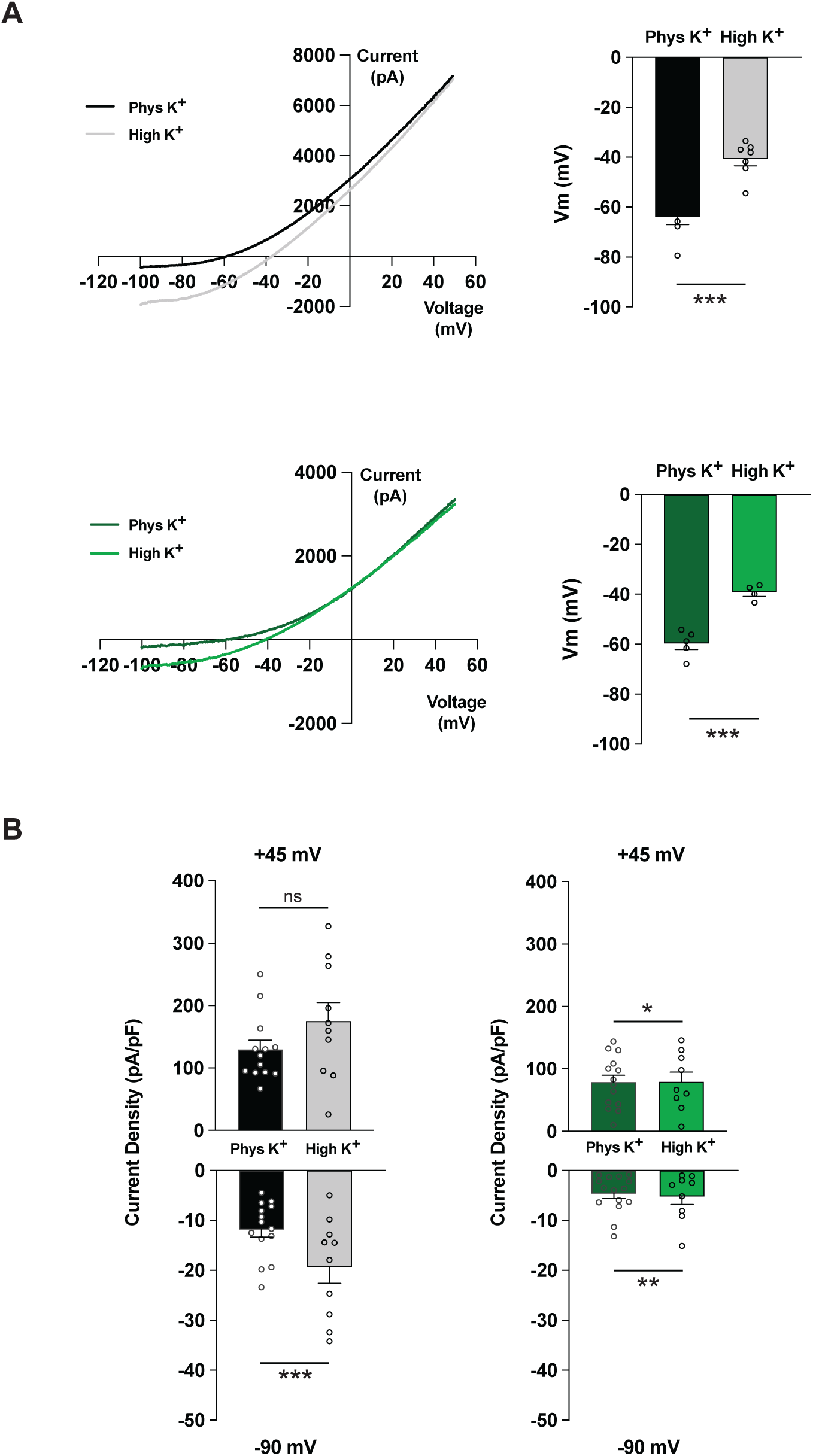
Potassium-dependent currents and reversal potentials of WT and c.1107del TRESK channels. **A.** Whole-cell currents were recorded using a voltage-ramp protocol (from −100 mV to +50 mV). Left: representative ramp-evoked current traces of both WT (grey) and c.1107del (green) under 5 mM (physiological) and 30 mM (elevated) extracellular K⁺ concentrations. Right: calculation of reversal potentials (E_rev_) for each cell, plotted as mean ± SEM at physiological and high [K^+^]_e_ and genotypes (n=7 WT in grey, n=4-5 c.1107del in green) (***P≤0.0001, unpaired Student’s t-test). **B.** Quantification of current densities (pA/pF) measured at −90 mV and +45 mV of WT (grey) and c.1107del (green) TRESK at both physiological and high [K^+^]_e_. Each data point represents an individual cell; bars indicate mean ± SEM (n=10-14 for WT and n=9-15 for c.1107del, ns non-significant P>0.05,*P≤0.05, **P≤0.01, unpaired Student’s t-test).

We next analyzed the voltage dependence of homomeric WT, c.1107del mutant, and heteromeric WT/mutant channels to assess how the c.1107del variant affects human TRESK activation. Normalized conductance–voltage (G/G_max_–Vm) relationships were fitted with a Boltzmann function to estimate the half-activation voltage (V₁_/_₂) and slope factor (k). The mean values from both parameters did not differ significantly among WT, c.1107del, and heteromeric WT/mutant channels showing that c.1107del does not affect the voltage sensitivity of TRESK activation (P=0.98 in WT vs. homomeric mutant and P=0.99 in WT vs. heteromeric WT/mutant, one-way ANOVA; Table 1). Overall, the c.1107del mutation preserves the fundamental biophysical properties of TRESK, including channel gating and ion selectivity.

**Table 1.**
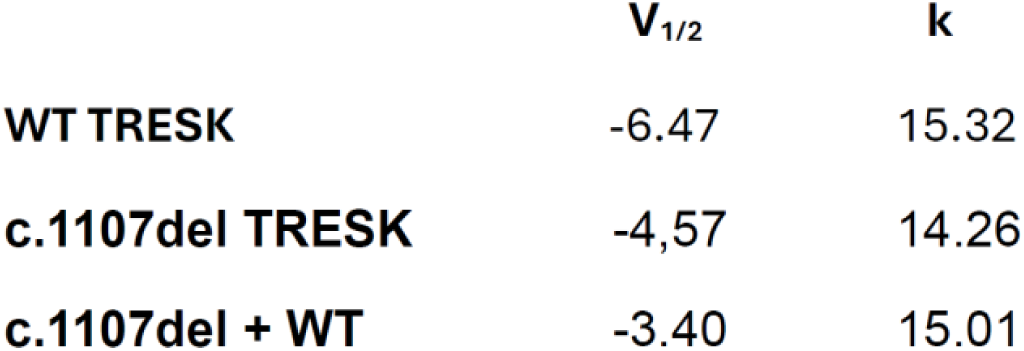
Voltage-dependence of activation for WT, c.1107del, and heterozygous TRESK channels. Normalized conductance–voltage (G/G_max_-Vm) relationships were obtained from step pulses (holding potential −60 mV, from −100 mV to +60 mV). Apparent conductance was calculated as G(V) = I(V)/(V − Erev) and normalized to G_max_. Data were fitted with a standard Boltzmann function to estimate V₁/₂ and slope factor (k). Points near E_rev_ were excluded to avoid numerical artifacts. Values are presented as mean ± SEM (n=270 WT, n=297 c.1107del). No significant differences were observed among WT, c.1107del, and heterozygous channels, indicating that c.1107del does not alter the voltage sensitivity of TRESK activation (P=0.98 for WT vs. homomeric mutant; P=0.99 for WT vs. heteromeric mutant, one-way ANOVA).

|  | **V<sub>1/2</sub>** | **k** |
| --- | --- | --- |
| **WT TRESK** | -6.47 | 15.32 |
| **c.1107del TRESK** | -4,57 | 14.26 |
| **c.1107del + WT** | -3.40 | 15.01 |

### The c.1107del TRESK variant forms functional heteromeric channels but shows reduced membrane trafficking

It has been described that some K2P channels can form heterodimeric channels with other members of the K2P family^36,37^. TRESK can form heterodimeric channels with TREK1 and TREK2, which have mixed properties^29^. Therefore, we next determined whether c.1107del alters TRESK heteromerization with other K2P subunits through single-molecule pulldown (SiMPull) assays. We quantified protein-protein interactions between EGFP-tagged WT and EGFP-c.1107del TRESK and HA-tagged TRESK and HA-TREK1. Co-pulldown events were detected as discrete fluorescent spots corresponding to heteromeric channel complexes at the single-molecule level (Fig. 5). Spot counts were normalized to the EGFP-TRESK - HA-TRESK condition to allow direct comparison across c.1107del and WT TRESK. As expected, TRESK-EGFP co-isolated efficiently with TRESK-HA (1.0 ± 0.02) and TREK1-HA (2.1 ± 0.2), above the background detected in TRESK-EGFP alone (0.2 ± 0.03). Consistenly, c.1107del-EGFP subunits displayed equivalent co-localization levels with both HA-TRESK and HA-TREK1 (1.1 ± 0.05 and 1.8 ± 0.1, respectively), again above its corresponding negative control (0.2 ± 0.04 spots) and not different from WT (P=0.22 for interactions with HA-TRESK and P=0.21 for interactions with HA-TREK1, n=10, unpaired Student’s t-test; Fig. 5). Hence, the c.1107del variant is still able to form homomeric and TREK1-containing heteromeric complexes, indicating that decreased currents seen electrophysiologically is not likely to be caused by defective interaction. We next compared the subcellular localization of WT and c.1107del variant by confocal microscopy to learn more about the mechanisms behind its partial loss of function. Quantitative colocalization using Mander’s coefficient showed a significantly lower overlap with the plasma membrane marker in c.1107del subunits (0.9 ± 2.8; n=12) compared to WT (1.8 ± 1.4; n=24, P < 0.001, unpaired t-test, Fig. 6).

**Fig 5.**
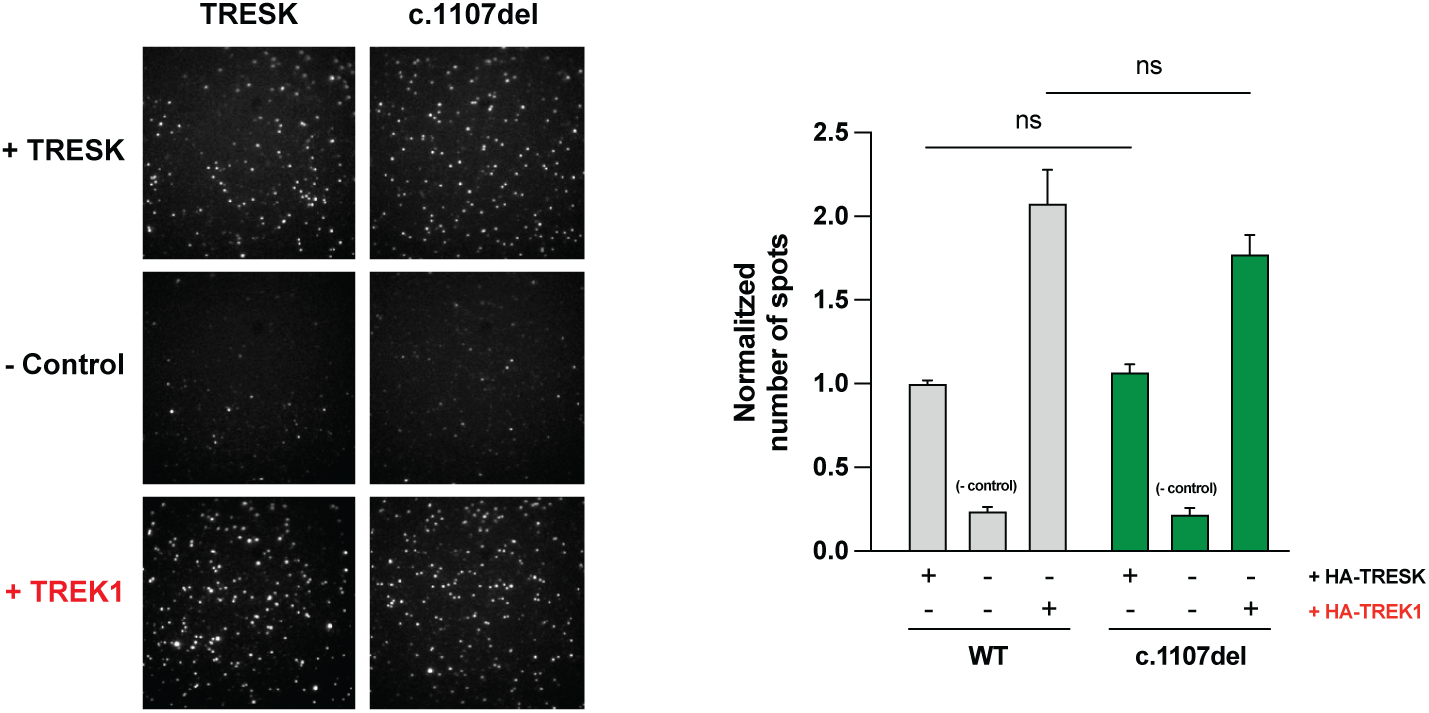
TRESK interaction with TRESK and TREK1 channel subunits (SimPull assay). Representative images showing that EGFP-hTRESK-WT (grey) and EGFP-hTRESK-c.1107del (green) are efficiently pulled down by both HA-TRESK and HA-TREK1. Immobilized anti-HA antibodies capture HA-tagged channels (TRESK and TREK1), and co-localization with GFP signal indicates subunit interaction. Quantification of the normalized number of GFP-positive spots pulled down by HA-TRESK or HA-TREK1. Both EGFP-hTRESK-WT and EGFP-hTRESK-c.1107del interact similarly with TRESK and TREK1, with no significant differences between WT and mutant. Data are mean ± SEM (n=10 Simpull experiments; ns non-significant P>0.05, unpaired Student’s t-test).

**Fig 6.**
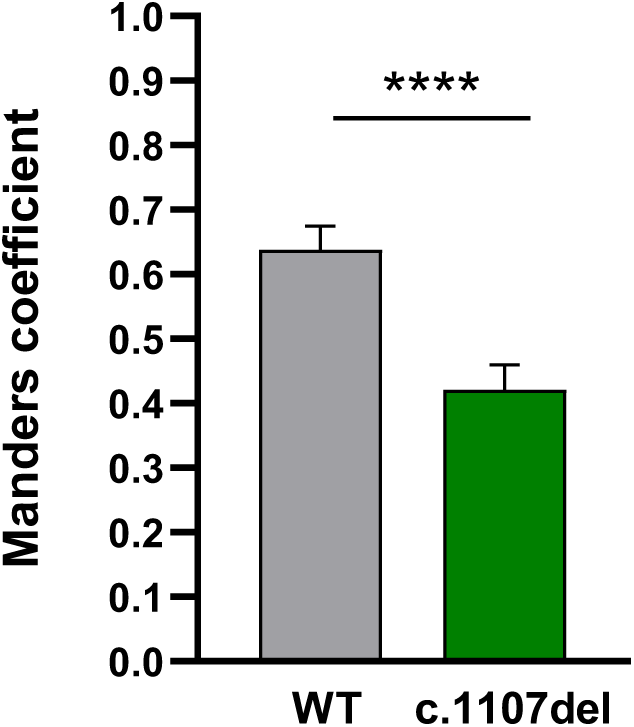
Plasma membrane expression of WT and c.1107del TRESK channels. Co-localization between fluorescently tagged TRESK constructs (WT-hTRESK-mCherry-C2 in red and c.1107del-hTRESK-EGFP-C2 in green), and Concanavalin A-Alexa 594 as the membrane marker reporting cell surface expression. Quantification using Mander’s overlap coefficient. c.1107del shows a significanty reduced co-localization with ConA compared to WT, indicating lower surface expression. Data are presented as arbitrary units (mean ± SEM) (n=270 for WT, n=297 for c.1107del, ****P < 0.0001, unpaired Student’s t-test).

### In silico analysis reveals altered phosphorylation pattern and decreased C-terminal hydrophobicity in the mutant TRESK

Considering that the c.1107del variant results in an elongated C-terminal domain, we investigated possible modifications to its post-translational regulation as well as biophysical characteristics using bioinformatic tools. Six new phosphorylation sites— three serines, two threonines, and one tyrosine—were identified by NetPhos analysis in the mutant C-terminus of TRESK, that may be targeted by distinct protein kinases including PKA and PKC. Conversely, there were no high-confidence phosphorylation sites predicted in the C-terminus of TRESK WT. In addition, the hydrophobic peak between residues 360 and 375 observed in the native sequence was significantly attenuated in the mutant protein, according to ProtScale analysis (data not shown), potentially modifying interactions of the C-terminal domain of the channel with membrane lipids.

## Discussion

TRESK, together with other K2P channels such as TREK1, TREK2 and TRAAK, provides the major background K⁺ conductances that stabilizes the resting membrane potential (RMP) and prevents excessive neuronal firing^2,7,8^. Although TRESK may contribute to the RMP, heterologous expression produces minor effects on resting potential of F-11 cells^9^ and TRESK deletion does not shift the RMP of sensory neurons^7^, likely due to compensatory K2P activity. Notably, neuronal excitability can change without any alteration in RMP due to the role of some K2P channels in the subthreshold range of the membrane potential. For instance, DRG neurons exhibit increased excitability after sciatic nerve axotomy despite significant changes in RMP were observed^8^. Beyond its debated contribution to the RMP, TRESK suppresses depolarising inputs and limits action potential initiation, acting as a key regulator on neuronal excitability. Consequently, TRESK appears to play an important role modulating responses to depolarizing stimuli^38^, as shown by its pharmacological activation with cloxyquin ^39,40^, which enhances currents and suppresses DRG neuronal excitability^28^, revealing its impact on sensory responsiveness.

Our findings reveal that the c.1107del (p.(Met370Cysfs)) frameshift mutation reduces TRESK-mediated background currents, consistent with a partial loss of function. Under heterozygous conditions, current densities fell between WT and mutant homozygous conditions, indicating a haploinsufficiency mechanism, whereby a single functional *KCNK18* allele is insufficient to maintain normal conductance. The reduction in TRESK K^+^ currents caused by the c.1107del mutation is expected to enhance nociceptor excitability, offering a plausible molecular contribution to the patient’s burning pain and paresthesias, due to TRESK expression in different subtypes of sensory neurons, including nociceptors. This phenotype aligns with previous studies associating *KCNK18* variants with altered excitability in migraine as well as other complex neurological disorders^27–29,41,42^. For instance, the F139WfsX24 frameshift mutation results in a complete loss of function with dominant-negative effect on other K2P channels^27,28^, while c.1107del results in a partial loss of function with no evidence of dominant-negative interference. Other reported substitutions identified in migraine patients —including R10G, S231P, and A233V—which impair TRESK current^29,41^ whereas W101R, also exhibiting markedly reduced currents, was found in a patient with intellectual disability and migraine with brainstem aura^31^, underscoring a variant-specific impact on TRESK function. The c.1107del *KCNK18* variant was identified in a patient with progressive SFN characterized by burning pain, sensory disturbances, and autonomic symptoms. This case illustrates the phenotypic heterogeneity associated with sensory neuropathies, in which symptom onset, severity, and autonomic involvement can vary widely even among carriers of pathogenic variants^43^. Such variability in clinical presentation highlights the complexity of linking genotype to clinical phenotype. Although our electrophysiology data cannot completely explain the patient’s symptoms, they offer a plausible molecular contribution to an altered sensory processing.

In addition, partial reduction in TRESK activity may influence neuronal excitability indirectly through interactions with other ion channels^29,44^. TRESK can form heteromeric complexes with TREK1/2 resulting in mixed K2P populations with distinct functional properties (e.g. calcium-mediated regulation of the heteromeric channel). It can also indirectly influence Na_v_ activation by RMP modulation^45,46^. SiMPull assays revealed that c.1107del form heteromers with TREK1 as efficiently as WT, generating equivalent oligomerization profiles. By comparison, the C110R variant, located adjacent to the selectivity filter and identified in migraineurs and controls, results in a complete loss-of-function while preserving TREK1/2 interactions and excitability in iPSC-derived nociceptors^41^. While channel assembly is preserved, we found that the number of c.1107del channels reaching the plasma membrane is substantially reduced, indicating defective trafficking to cell surface as the main deficit. Electrophysiological analysis further confirmed that the mutation does not affect intrinsic channel gating behavior. Although TRESK is not voltage-gated, apparent voltage dependence under asymmetric physiological solutions reflects changes in driving force and intrinsic permeation, rather than classical gating. Thus, we measured normalized conductance using (G/G_max_)–voltage relationships and Boltzmann fits to provide cautious comparison of TRESK activation. WT and mutant channels displayed comparable V₁_/_₂ values and k factors, indicating conserved voltage sensitivity. Despite K2P channels were initially described as voltage-insensitive as they do not possess a canonical voltage-sensing domain, other studies reported that they can be voltage gated through a ion check valve mechanism^47^. Likewise, ionic-selectivity measurements reflects that both WT and c.1107del TRESK reproduce the differential impact of high [K⁺]_e_ on the electrochemical driving force, substantially enhancing K⁺ entry more than K⁺ efflux^48^, indicating the ion-permeation mechanism remains intact despite reduced current amplitude.

The location of the c.1107del mutation in the distal C-terminal domain suggests additional regulatory consequences. The C-terminus plays essential dual roles in TRESK activity. The proximal region, including basic residues within the KLVQNR motif, contributes to gating modulation and transmits regulatory inputs. Interestingly, modifications in this region can lock the channel in a low-activity, calcineurin-insensitive state^49^. On the other hand, the distal C-terminus-were c.1107del occurs-contains a short hydrophobic segment (VMLFFA) that is predicted to interact with the inner leaflet of the plasma membrane, based on its strong hydrophobicity. Progressive deletions of this hydrophobic patch strongly reduce or abolish TRESK currents, while replacements with more membrane-preferring hydrophobic sequences enhance channel activity. It confirms that anchoring of the C-terminus distal tail promotes TRESK function^49^. The c.1107del mutation replaces the hydrophobic VMLFFA motif with a more polar set of residues (VCYSLQ), shifting the chemical character of the C-terminus. This change—from non-polar, membrane-anchoring residues to a polar, uncharged amino acids—likely lessen insertion into the lipid core and compromises coupling to the membrane surface. In addition, the C-terminus domain in TRESK and TREK1 likely contributes to dimer formation and channel trafficking to the membrane, as partial or total deletion of the C-terminal domain provokes a retention of the channel in the endoplasmic reticulum and avoids the formation of functional channels in the membrane^29,34,50^. Besides, the new C-terminal sequence introduces multiple high-confidence phosphorylation sites, according to bioinformatic analysis (NetPhos), including PKA-predicted Ser23 and PKC-predicted Thr25/Thr26. Phosphorylation of these residues may cause additional downstream effects and post-translational regulation. PKA is known to inhibit TRESK activity via phosphorylation of Ser252 in the intracellular loop^51^. New PKA/PKC-sensitive sites may therefore result in synergistic or additive inhibitory effects.

Taken together, the combined features linked to c.1107del—including partial loss of function, reduced surface expression, maintained K^+^ selectivity and channel oligomerization, and domain-specific remodeling of the C-terminus—outline a complex pattern of dysfunction. These molecular changes offer a potential mechanistic pathway to explain a lower activation threshold of sensory neurons contributing to patients’ symptoms. Future research will be required to define the physiological impact of c.1107del on neuronal excitability and clinical heterogeneity.

